# Roles of Lipopolysaccharide Glycosyltransferases in Maintenance of *Helicobacter pylori* Morphology, Cell Wall Permeability, and Antimicrobial Susceptibilities

**DOI:** 10.1101/2023.05.25.542268

**Authors:** Xiaoqiong Tang, Tiankuo Yang, Yalin Shen, Xiaona Song, Mohammed Benghezal, Barry J. Marshall, Hong Tang, Hong Li

**Affiliations:** West China Marshall Research Center for Infectious Diseases, Center of Infectious Diseases, West China Hospital, Sichuan University, Chengdu, China; Division of Infectious Diseases, State Key Laboratory of Biotherapy and Center of Infectious Diseases, West China Hospital, Sichuan University, Chengdu, China; Aviation Medical Appraisal Center, Civil Aviation Flight University of China, Guanghan, China; Helicobacter pylori Research Laboratory, School of Biomedical Sciences, Marshall Centre for Infectious Disease Research and Training, University of Western Australia, Nedlands, Australia; School of Biomedical Engineering, Marshall Laboratory of Biomedical Engineering, Shenzhen University Health Science Center, Shenzhen, China

**Keywords:** *Helicobacter pylori*, lipopolysaccharide, glycosyltransferases, morphology, antimicrobial susceptibilities

## Abstract

*Helicobacter pylori* unique lipopolysaccharide structure is essential in maintaining the cell envelop integrity and renders the bacterium natural resistance to cationic antimicrobial peptides (CAMPs). Our group has recently elucidated the complete set of LPS glycosyltransferase genes in *H. pylori* reference strain G27. Here, with a series of 8 systematically constructed LPS glycosyltransferase gene mutants (G27Δ*HP1578*, G27Δ*HP1283*, G27Δ*HP0159*, G27Δ*HP0479*, G27Δ*HP0102*, G27Δ*wecA*, G27Δ*HP1284* and G27Δ*HP1191*), we investigated the roles of *H. pylori* LPS glycosyltransferases in maintenance of cell morphology, cell wall permeability, and antimicrobial susceptibilities. We demonstrated that deletion of these LPS glycosyltransferase genes did not interfere with bacterial cell wall permeability, but resulted in significant morphological changes (coccoid, coiled “c”-shape, and irregular shapes) after 48 h growth as compared to the rod-like cell shape of the wild-type strain. Moreover, as compared with the wild-type, none of the LPS mutants had altered susceptibility against clarithromycin, levofloxacin, amoxicillin, tetracycline, and metronidazole. However, the deletion of the conserved LPS glycosyltransferases, especially the O-antigen initiating enzyme WecA displayed a dramatic increase in susceptibility to the CAMP polymyxin B and rifampicin. Taken together, our findings suggest that the LPS glycosyltransferases play critical roles in the maintenance of the typical spiral morphology of *H. pylori*, as well as resistance to CAMPs and rifampicin. The LPS glycosyltransferases could be promising targets for developing novel anti-*H. pylori* drugs.

**Importance:** *H. pylori* typical helical morphology, cell wall integrity, as well as resistance to cationic CAMPs and antimicrobials are significant factors for its long-term colonization and persistent infection in human gastric mucosa. Our results show that each of the 8 LPS glycosyltransferase genes (*HP1578*, *HP1283*, *HP0159*, *HP0479*, *HP0102*, *wecA*, *HP1284* and *HP1191*) deletion did not interfere with bacterial cell wall permeability, but resulted in significant loss of *H. pylori* typical helical shape. Furthermore, deletion of the conserved LPS glycosyltransferases, especially the O-antigen initiating enzyme WecA displayed a dramatic increase in susceptibility to the CAMP polymyxin B and rifampicin. Taken together, we believe that the LPS glycosyltransferases are good targets for developing novel anti-*H. pylori* drugs.

## Introduction

*Helicobacter pylori* infection is an infectious disease, infecting approximately half of the world’s population.^1^ Usually acquired in childhood, *H. pylori* infection is chronic and persistent, which can lead to the development of peptic ulcer, chronic gastritis, gastric cancer, and mucosa associated lymphatic tissue (MALT) lymphoma.^2^ *H. pylori* infection is the most important infectious cause of cancer worldwide, accounting for approximately 90% new cases of the global gastric cancer.^3^ Thus, *H. pylori* eradication has been recommended as the primary strategy for the prevention of gastric cancer development.^4–6^

Like other Gram-negative bacteria, the surface of *H. pylori* is composed of an asymmetric outer membrane with phospholipids in the inner leaflet and lipopolysaccharide (LPS) exclusively anchored in the outer leaflet.^7, 8^ As a major constituent of the outer membrane, LPS plays an essential role in maintaining the cell envelop integrity and forming an effective barrier that is impermeable to many toxic compounds, including antibiotics.^9^ *H. pylori* LPS is composed of three domains: 1) the hydrophobic lipid A anchoring LPS in the outer membrane; 2) the central core oligosaccharide; and 3) the distal O-antigen.^8^ *H. pylori* LPS lipid A is constitutively modified through dephosphorylation and deacylation to confer *H. pylori* intrinsic resistance to cationic antimicrobial peptides (CAMPs), and the ability to evade Toll-like receptor 4 (TLR-4) recognition.^10^ In addition, the molecular mimicry between *H. pylori* LPS O-antigen and host Lewis blood group antigens also helps camouflaging *H. pylori* from detection by host immune surveillance.^11^ Thus, the unique *H. pylori* LPS structure plays a critical role in the establishment of *H. pylori* colonization and persistent infection within the host gastric niche.

Our group has recently elucidated the complete LPS structure in *H. pylori* reference strain G27^12^ (**Fig. 1**). Unlike the core oligosaccharide of LPS in other Gram-negative bacteria having an inner and outer core, we demonstrated that *H. pylori* LPS core oligosaccharide is a short and conserved hexa-saccharide (-Glc-Gal-DD-Hep III-LD-Hep II-LD-Hep I-KDO-). We identified HP1284 as the Hep III glycosyltransferase,^12^ while the Hep I and Hep II glycosyltransferases have been previously identified as HP0279 and HP1191, respectively.^13, 14^ *H. pylori* LPS O-antigen was previously proposed to contain Lewis antigen only.^15^ However, through systematic mutagenesis of glycosyltransferase genes in strain G27 combined with LPS structural analysis, we demonstrated that *H. pylori* LPS O-antigen is a long and linear structure encompassing a trisaccharide (-DD-Hep-Fuc-GlcNAc-) termed as Trio, a glucan, a DD-heptan, and the terminal Lewis antigens. Of note, the Trio, glucan, and heptan were previously assigned as the outer core domain.^16^ We also identified HP0102 as the Trio Fuc glycosyltransferase, HP1283 as the heptan glycosyltransferase, and HP1578 as the GlcNAc glycosyltransferase responsible for initiating Lewis antigen synthesis onto the heptan.^17^ Our study also enabled the assignment of the GlcNAc in the Trio as the first sugar of the long O-antigen, and the GlcNAc in the Trio is added by the O-antigen initiating enzyme WecA (HP1581).^17^ The DD-Hep in the Trio and the glucan have been previously reported to be transferred by HP0479^13^ and HP0159,^18^ respectively.

**Fig. 1.**
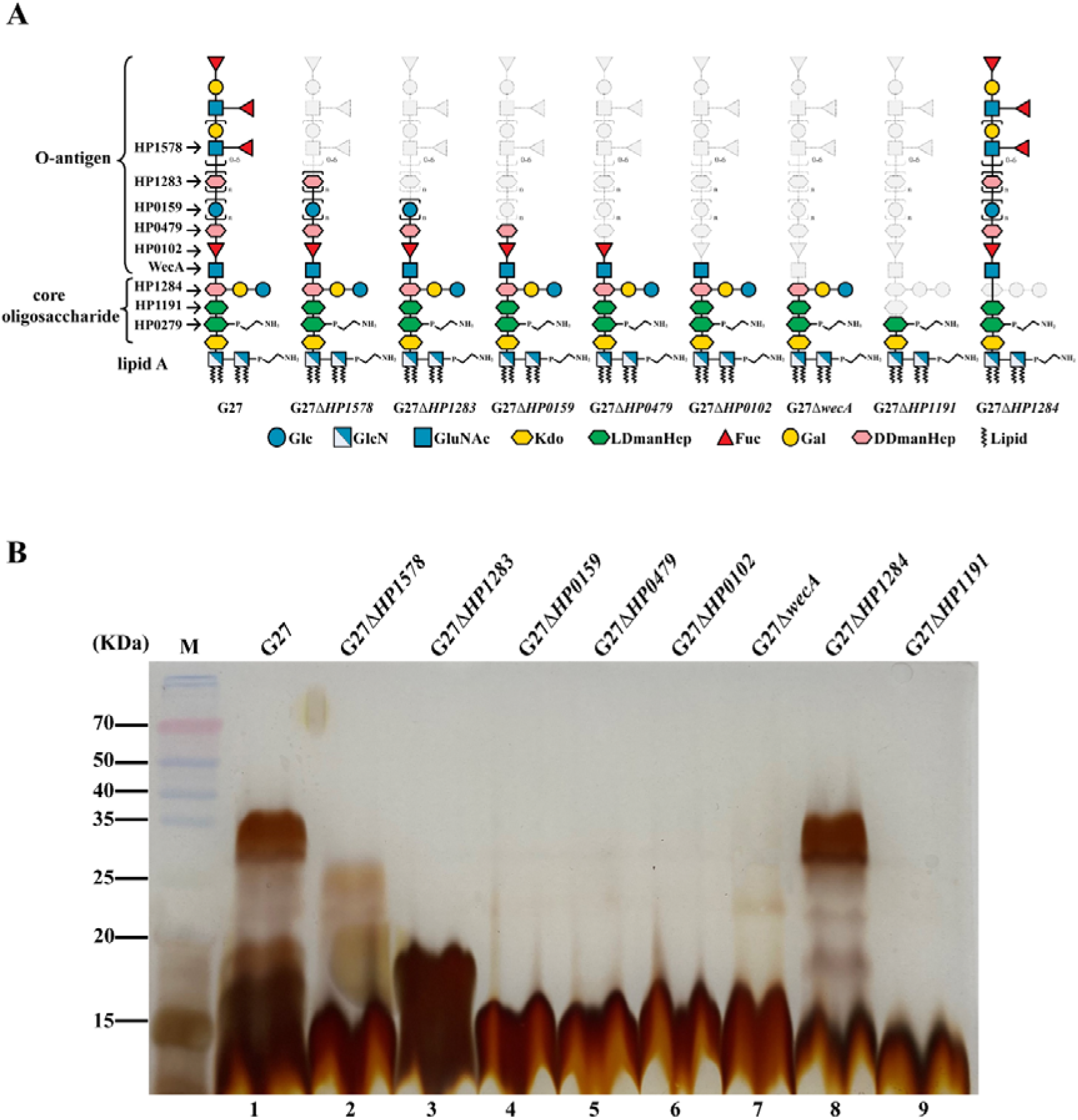
LPS structural characterization of *H. pylori* LPS mutants. (A) Chemical structure model of wild-type G27 and its associated LPS mutants. The gene was named according to standard *H. pylori* strain 26695. The gray represents the truncated portion of LPS. Gal, galactose; Glc, glucose; Fuc, fucose; GlcNAc, nitroacetyl glucosamine; DDmanHep, D-glycerol-D-mannoheptanose; LDmanHep, L-glycerol-D-mannoheptanose; GlcN, glucosamine; Kdo, 3-deoxy-D-mannose-octanoic acid. (B) LPS structural characterization of wild-type G27 and its associated LPS mutants detected by silver staining. M, molecular weight marker.

Considering the identification of the complete set of LPS glycosyltransferase genes in *H. pylori* strain G27, and in view of the essential roles played by *H. pylori* LPS in bacterial survival and host-pathogen interactions, the purpose of this study was to characterize the underlying changes in bacterial fitness, cell morphology, cell wall permeability, and antimicrobial susceptibility among a series of systematically constructed LPS glycosyltransferase gene mutants in G27. Our data showed that deletion of these LPS glycosyltransferase genes dose not interfere with bacterial fitness and cell wall permeability, but affects the spiral and rod-like morphology. Moreover, antibiotic susceptibility testing revealed that deletion of LPS glycosyltransferase genes increases *H. pylori* sensitivity to polymyxin B and rifampicin. These findings suggest that the LPS glycosyltransferases play critical roles in the maintenance of the typical spiral morphology of *H. pylori*, as well as resistance to antibiotics.

## Results

### Deletion of LPS glycosyltransferase genes leads to various degree of *H. pylori* LPS truncation, but does not interfere with bacterial fitness

A panel of 8 LPS glycotransferase gene mutants constructed previously in the same background strain G27 was included in this study (**Table 1**). Of note, numerous attempts to delete *HP0279* gene coding for the Hep I transferase was unsuccessful in our laboratory, which is consistent with a previous study,^19^ suggesting that HP0279 is essential in *H. pylori*. The LPS structures of the G27 wild-type strain and the 8 isogenic mutants have been previously elucidated and summarized in **Fig. 1A**.^17^ Their LPS profiles were compared in this study by silver staining on SDS-PAGE gel (**Fig. 1B**). As expected, apparent LPS truncation was observed in 7 of the 8 mutants: G27Δ*HP1578*, lacking the distal Lewis antigen only; G27Δ*HP1283*, lacking DD-heptan and the the Lewis antigen; G27Δ*HP0159*, lacking glucan, the DD-heptan, and the Lewis antigen; G27Δ*HP0479*, lacking Hep residue of the Trio, the glucan, the DD-heptan, and the Lewis antigen; G27Δ*HP0102*, lacking fuc and Hep residues of the Trio, the glucan, the DD-heptan, and the Lewis antigen; G27Δ*wecA*, lacking the whole O-antigen; and G27Δ*HP1191*, lacking Hep II, Hep I resides and the whole O-antigen. As for G27Δ*HP1284*, it’s LPS profile was similar to that of the wild-type strain (**Fig. 1B, Lane 8**), as it has been previously demonstrated that the deletion of the Hep III transferase gene *HP1284* led to the loss of the Hep III residue and the adjoining Glc-Gal residues only, whereas the O-antigen ligation was not affected, which was attached onto Hep II residue of the incomplete core-oligosaccharide.^12^ ^17^

**Table 1.** Bacterial strains and plasmids used in this study.

| Strain or plasmid name | Description | LPS biosynthesis | Reference or source |
| --- | --- | --- | --- |
| ***H. pylori* strains** |  |  |  |
| G27 | Wild-type strain, streptomycin resistant | Full-length LPS | Li et al., 2019 |
| G27ΔHP1578 | Clean deletion of *HPG27_1515* in G27 | Deletion of Lewis antigen | Li et al., 2019 |
| G27ΔHP1283 | Clean deletion of *HPG27_1235* in G27 | Deletion of Lewis antigen and heptan | Li et al., 2019 |
| G27ΔHP0159 | Clean deletion of *HPG27_146* in G27 | Deletion of Lewis antigen, DD-heptan, and glucan | Li et al., 2019 |
| G27ΔHP0479 | Clean deletion of *HPG27_437* in G27 | Deletion of Lewis antigen, DD-heptan, glucan, and LD-heptose of Trio | Li et al., 2019 |
| G27ΔHP0102 | Clean deletion of *HPG27_94* in G27 | Deletion of Lewis antigen, DD-heptan, glucan, Trio LD-heptose and fucose | Li et al., 2019 |
| G27ΔwecA | Clean deletion of *HPG27_1518* in G27 | Deletion of the entire O-antigen | Li et al., 2019 |
| G27ΔHP1284 | Clean deletion of *HPG27_1236* in G27 | Deletion of side-chain and heptose III in the core oligosaccharide | Li et al., 2019 |
| G27ΔHP1191 | Clean deletion of *HPG27_1136* in G27 | Deletion of the entire O-antigen, and side-chain, heptose III and heptose II in the core oligosaccharide | Li et al., 2019 |
| 26695 | Wild-type strain, streptomycin resistant | Full-length LPS | Stored in our lab |
| 26695ΔwecA | Clean deletion of *HP1581* in 26695 | Deletion of the entire O-antigen | This work |
| J99 | Wild-type strain | Full-length LPS | Stored in our lab |
| J99ΔwecA | Deletion of *wecA* in J99 by replacing it with *difH-rpsI-cat-difH* | Deletion of the entire O-antigen | This work |
| P12 | Wild-type strain | Full-length LPS | Stored in our lab |
| P12Δ*wecA* | Deletion of *wecA* in P12 by replacing it with *difH-rpsL-cat-difH* | Deletion of the entire O-antigen | This work |
| **Plasmids** |  |  |  |
| pWecA-AB-difH-RC | pGEM<sup>®</sup>-T Easy vector containing sequences flanking *HPG27_1518*, with *difH-rpsL-cat-difH* inserted between the flanking sequences |  | Li et al., 2019 |

To investigate whether deletion of LPS glycosyltransferase genes or LPS truncation interfere with bacterial fitness, growth curves of the G27 wild-type and the 8 LPS mutants were performed in liquid broth, and the optical densities were measured every 6 or 12 hours for up to 48 h. Interestingly, all the 8 LPS mutants displayed similar or higher growth rate than that of the wild-type strain up to 36 h, suggesting that the deletion of LPS glycosyltransferase genes do not negatively impact bacterial fitness. Of note, G27Δ*wecA* mutant had the most robust growth among all the stains up to 36 h. From 36 h to 48 h, the optical densities of G27Δ*wecA*, G27Δ*HP0102*, G27Δ*HP1284*, G27Δ*HP1283*, and G27Δ*HP0479* still increased; whereas the G27 wild-type, G27Δ*HP1578*, and G27Δ*HP1191* strains had a marked decrease, suggesting that these three strains had reached the decline phase (**Fig. 2**).

**Fig. 2.**
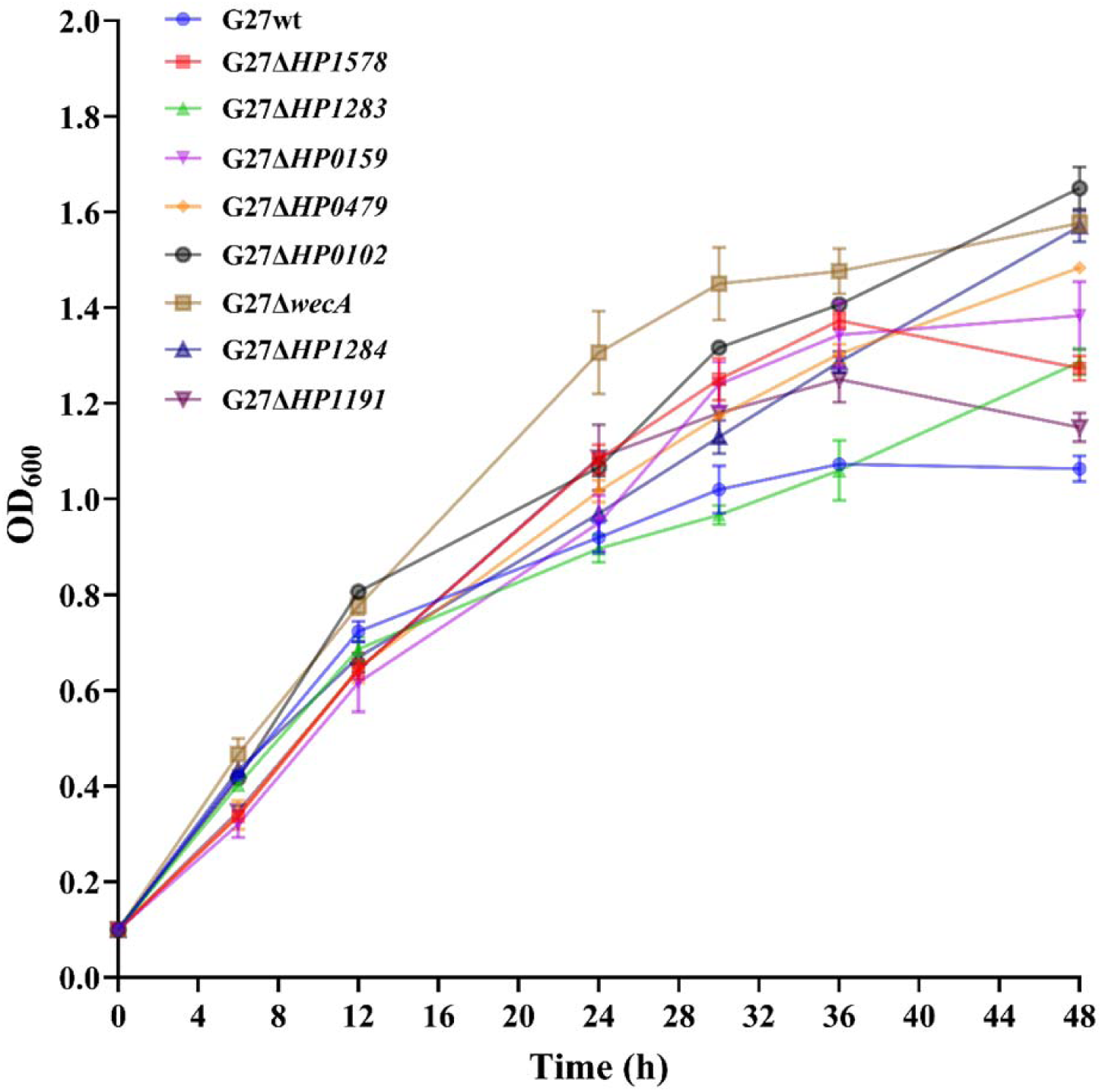
Deletion of *H. pylori* LPS glycosyltransferase genes does not interfere with bacterial fitness. Growth curve analysis of wild-type G27 and LPS mutants through measuring the optical density at 600 nm (OD_600_) wavelength of the cultures every 6 h and/or 12 h for up to 48 h. Each strain was grown in triplicate, and data from two independent experiments are represented as mean with standard error of the mean (± SEM).

### Deletion of LPS glycosyltransferase genes affects the spiral and rod-like morphology

As LPS is one of the major constituents of the outer membrane, the perturbation of LPS biosynthesis upon the deletion of corresponding glycosyltransferase genes may have a great effect on the cell envelope, and thus causing morphological changes to the helical shape of *H. pylori*.

To investigate the roles played by LPS glycosyltransferase genes on *H. pylori* morphologies, the G27 wild-type and the 8 LPS mutants were visualized by gram-staining after 24 h and 48 h of culture. All the strains exhibited the characteristic helical and rod-like cell shape after 24 of culture (**Fig. 3A**). However, compared with the wild-type strain, apparent elongated cell shape was observed in G27Δ*HP0479*, G27Δ*wecA*, and G27Δ*HP1284* mutants. Interestingly, after 48 of culture, the rod-like cell shape was still clearly observed in wild-type, G27Δ*HP0159*, G27Δ*HP0479*, and G27Δ*HP1191,* whereas the cells in G27Δ*HP1578* and G27Δ*wecA* were almost completely coccoid, and a mixture of coccoid, coiled “c”-shape, and irregular shapes were observed in G27Δ*HP1283*, G27Δ*HP0102*, and G27Δ*HP1284* (**Fig. 3B**).

**Fig. 3.**
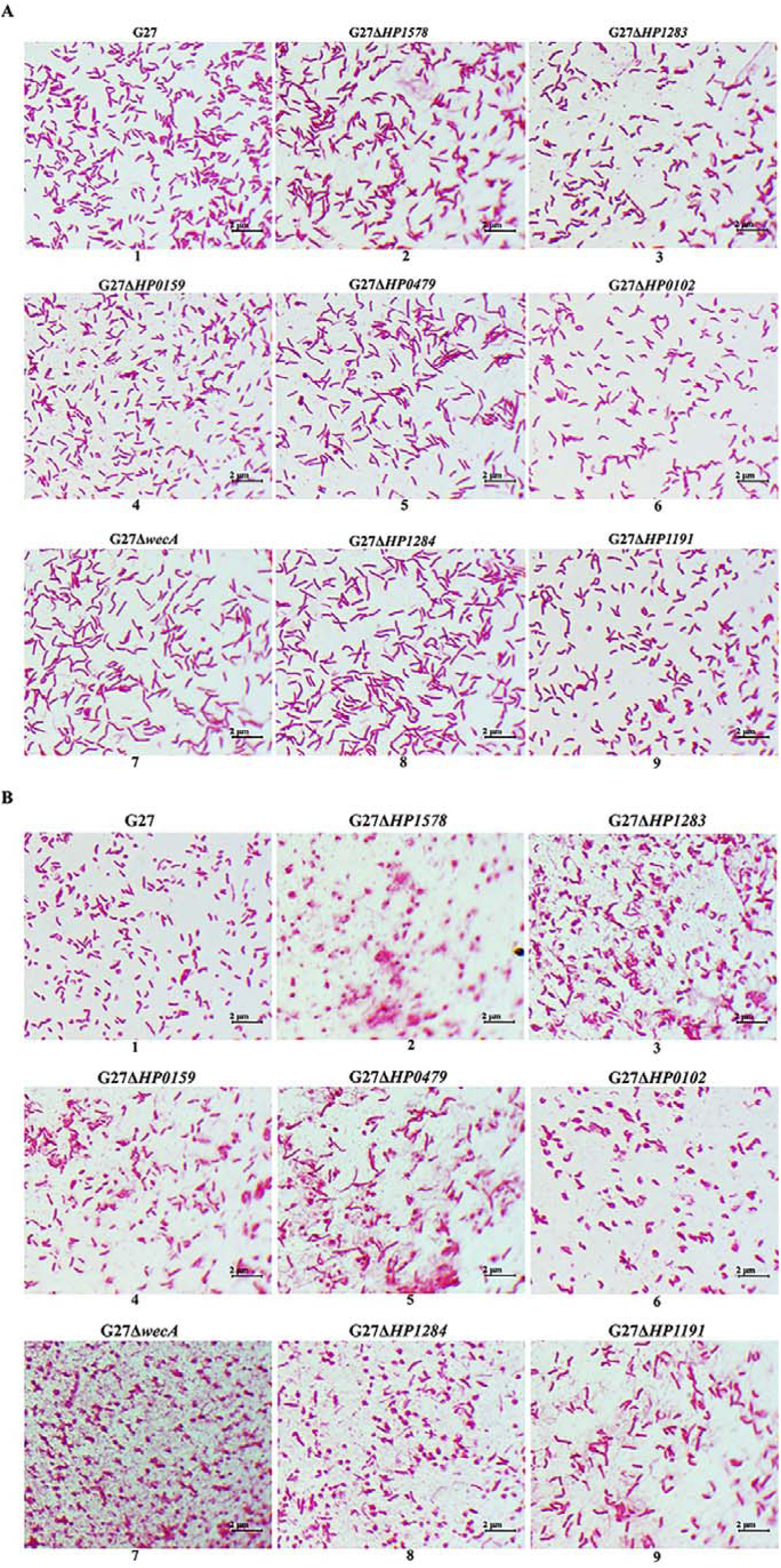
*H. pylori* cell morphology is perturbed by deletion of LPS glycosyltransferase genes. Gram-staining depicting morphological patterns of *H. pylori* G27 wild-type and LPS mutants after 24h (A) and 48h (B) growth. Images were taken using Light microscope (1000x).

### Deletion of LPS glycosyltransferase genes affects *H. pylori* sensitivity to polymyxin B

CAMPs are positively charged, which can interact with the negatively charged surface structures (primarily LPS lipid A) of most Gram-negative bacteria, inducing bacterial lysis and death.^20^ Thus, CAMPs represent an important component of the host innate immune system. Due to constitutive modification of LPS, *H. pylori* is naturally resistant to polymyxin B,^10^ the surrogate of host CAMPs in laboratory settings. Modification or variation in LPS structure has been reported to affect *H. pylori* resistance to polymyxin B.^12, 21, 22^ However, the roles played by LPS glycosyltransferase genes on *H. pylori*’s resistance to the CAMP polymyxin B has not been systematically studied.

Using the “gold standard” agar dilution method, MIC of polymyxin B was first determined in G27 wild-type and the 8 isogenic LPS mutants. The wild-type G27 had a polymyxin B MIC of 18.67 ± 7.05 μg/mL, and both of the G27Δ*HP1578* and G27Δ*HP1283* mutants had a MIC of 13.33 ± 2.67 μg/mL, which was comparable to the MIC of the wild-type strain. The polymyxin B MICs in G27Δ*HP0159,* G27Δ*HP479,* G27Δ*HP0102*, *HP1284*, and G27Δ*HP1191* were 2.00 ± 0.00, 1.50 ± 0.50, 6.67 ± 1.33, 3.25 ± 2.37, and 2.33 ± 0.88 μg/mL, respectively, which was a marked decrease in resistance to polymyxin B as compared to the wild-type (**Table 2**, **Fig. 4A-B**).

**Fig. 4.**
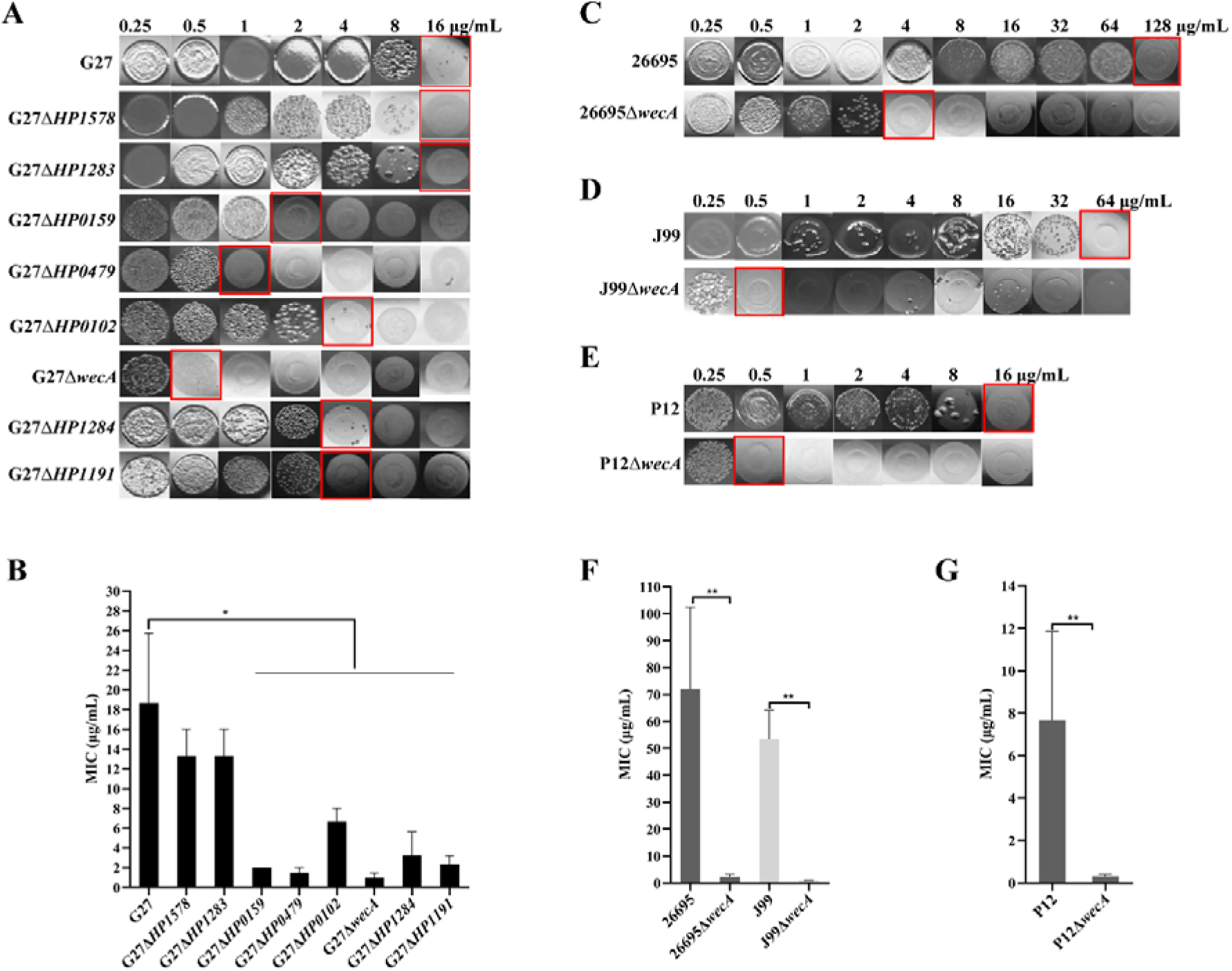
Deletion of LPS glycosyltransferase genes leads to reduction of *H. pylori* resistance to polymyxin B. Resistance patterns of *H. pylori* G27 (A), 26695 (C), J99 (D), P12 (E) and their associated LPS mutants to polymyxin B were profiled by agar dilution assay. The strains were grown in brucella broth infusion (BHI) blood agar plates supplemented with various concentrations of polymyxin B. The minimum inhibitory concentration (MIC), outlined in red, was defined as the lowest concentration of antibiotics at which bacterial grew was obviously inhibited. (B) MICs of wild-type G27 and its LPS mutants to polymyxin B obtained by three independent experiments. (F-G) MICs of wild-type 26695, J99 and P12 and their Δ*wecA* mutants to polymyxin B obtained by three independent experiments. All values were shown as mean with standard error of the mean (± SEM). *, *P* < 0.05, **, *P* < 0.01, ***, *P* < 0.001; Student’s t-test, two-tailed.

**Table 2.** Minimum inhibitory concentrations (MICs) of *H. pylori* wild-type and LPS glycosyltransferase genes mutant strains against antibiotics.

| Strains | Polymyxin B<br>($\mu\text{g/mL}$) | Rifampicin<br>($\mu\text{g/mL}$) | Clarithromycin<br>($\mu\text{g/mL}$) | Levofloxacin<br>($\mu\text{g/mL}$) | Amoxicillin<br>($\mu\text{g/mL}$) | Tetracycline<br>($\mu\text{g/mL}$) | Metronidazole<br>($\mu\text{g/mL}$) |
| --- | --- | --- | --- | --- | --- | --- | --- |
| G27 | 18.67 ± 7.05 | 6.67 ± 1.33 | 0.03 ± 0.01 | 0.23 ± 0.02 | 0.04 ± 0.01 | 0.27 ± 0.05 | 3.30 ± 0.33 |
| G27ΔHP1578 | 13.33 ± 2.67 | 1.83 ± 0.16 | 0.03 ± 0.01 | 0.18 ± 0.04 | 0.04 ± 0.01 | 0.27 ± 0.05 | 3.30 ± 0.33 |
| G27ΔHP1283 | 13.33 ± 2.67 | 1.42 ± 0.36 | 0.03 ± 0.01 | 0.21 ± 0.02 | 0.03 ± 0.01 | 0.27 ± 0.05 | 3.30 ± 0.33 |
| G27ΔHP0159 | 2.00 ± 0.00 | 2.50 ± 0.76 | 0.02 ± 0.01 | 0.19 ± 0.04 | 0.04 ± 0.01 | 0.23 ± 0.02 | 3.00 ± 0.57 |
| G27ΔHP0479 | 1.50 ± 0.50 | 0.83 ± 0.08 | 0.02 ± 0.01 | 0.17 ± 0.02 | 0.03 ± 0.00 | 0.27 ± 0.05 | 3.00 ± 0.57 |
| G27ΔHP0102 | 6.67 ± 1.33 | 0.66 ± 0.08 | 0.03 ± 0.01 | 0.17 ± 0.02 | 0.04 ± 0.01 | 0.27 ± 0.05 | 3.33 ± 0.33 |
| G27ΔwecA | 0.96 ± 0.52 | 0.42 ± 0.04 | 0.02 ± 0.01 | 0.21 ± 0.02 | 0.03 ± 0.00 | 0.27 ± 0.05 | 3.33 ± 0.66 |
| G27ΔHP1284 | 3.25 ± 2.37 | 0.58 ± 0.08 | 0.02 ± 0.01 | 0.19 ± 0.04 | 0.04 ± 0.01 | 0.27 ± 0.05 | 3.17 ± 0.83 |
| G27ΔHP1191 | 2.33 ± 0.88 | 0.58 ± 0.08 | 0.03 ± 0.01 | 0.17 ± 0.02 | 0.04 ± 0.01 | 0.27 ± 0.05 | 2.50 ± 0.76 |
| 26695 | 72.00 ± 30.29 | 5.33 ± 1.33 | 0.08 ± 0.02 | 0.33 ± 0.08 | 0.08 ± 0.02 | 0.21 ± 0.04 | 85.3 ± 21.3 |
| 26695ΔwecA | 2.25 ± 0.94 | 1.08 ± 0.51 | 0.08 ± 0.02 | 0.31 ± 0.09 | 0.08 ± 0.02 | 0.21 ± 0.04 | 85.3 ± 21.3 |
| J99 | 53.33 ± 10.67 | 5.33 ± 1.33 | 0.02 ± 0.01 | 0.09 ± 0.03 | 0.03 ± 0.01 | 0.10 ± 0.04 | 0.67 ± 0.17 |
| J99ΔwecA | 0.83 ± 0.17 | 2.33 ± 0.88 | 0.02 ± 0.01 | 0.09 ± 0.02 | 0.03 ± 0.01 | 0.09 ± 0.02 | 0.63 ± 0.19 |
| P12 | 7.67 ± 4.18 | 6.00 ± 1.15 | 0.03 ± 0.01 | 0.23 ± 0.02 | 0.02 ± 0.00 | 0.21 ± 0.02 | 0.83 ± 0.17 |
| P12ΔwecA | 0.31 ± 0.09 | 2.16 ± 0.93 | 0.02 ± 0.01 | 0.21 ± 0.04 | 0.02 ± 0.00 | 0.23 ± 0.02 | 0.83 ± 0.17 |

Notably, G27Δ*wecA* had the lowest MIC of 0.96 ± 0.52 μg/mL, suggesting that *wecA* gene may play an important role in *H. pylori* resistance to polymyxin B (**Table 2**, **Fig. 4A-B**). To confirm the marked decrease of polymyxin B resistance in Δ*wecA* mutant is not strain-specific, we constructed Δ*wecA* mutant in another 3 different *H. pylori* strains: 26695Δ*wecA*, J99Δ*wecA*, and P12Δ*wecA*. The deletion of *wecA* in these 3 strains was confirmed by silver staining showing the loss of the whole O-antigen (**Fig. S1**). The polymyxin B MICs in wild-type 26695, J99, and P12 were 72.00 ± 30.29, 53.33 ± 10.67, and 7.67 ± 4.18, respectively. In comparison, the polymyxin B MICs in their Δ*wecA* mutants showed a marked 36, 63, and 25-fold decrease to 2.25 ± 0.94, 0.83 ± 0.17, and 0.31 ± 0.09, respectively (**Table 2**, **Fig. 4C-G**).

### Deletion of LPS glycosyltransferase genes does not affect *H. pylori* outer membrane permeability, but increases bacteria sensitivity to rifampicin

Given that LPS is the major component of *H. pylori* outer membrane, we investigated whether the deletion of LPS glycosyltransferase genes or LPS truncation had any effect on the outer membrane permeability. We measured the influx of fluorescent probe NPN among wild-type and the LPS mutants, and no NPN fluorescence intensity difference was observed (**Fig. 5**), suggesting that the deletion of the LPS glycosyltransferase genes does not affect *H. pylori* outer membrane permeability.

**Fig. 5.**
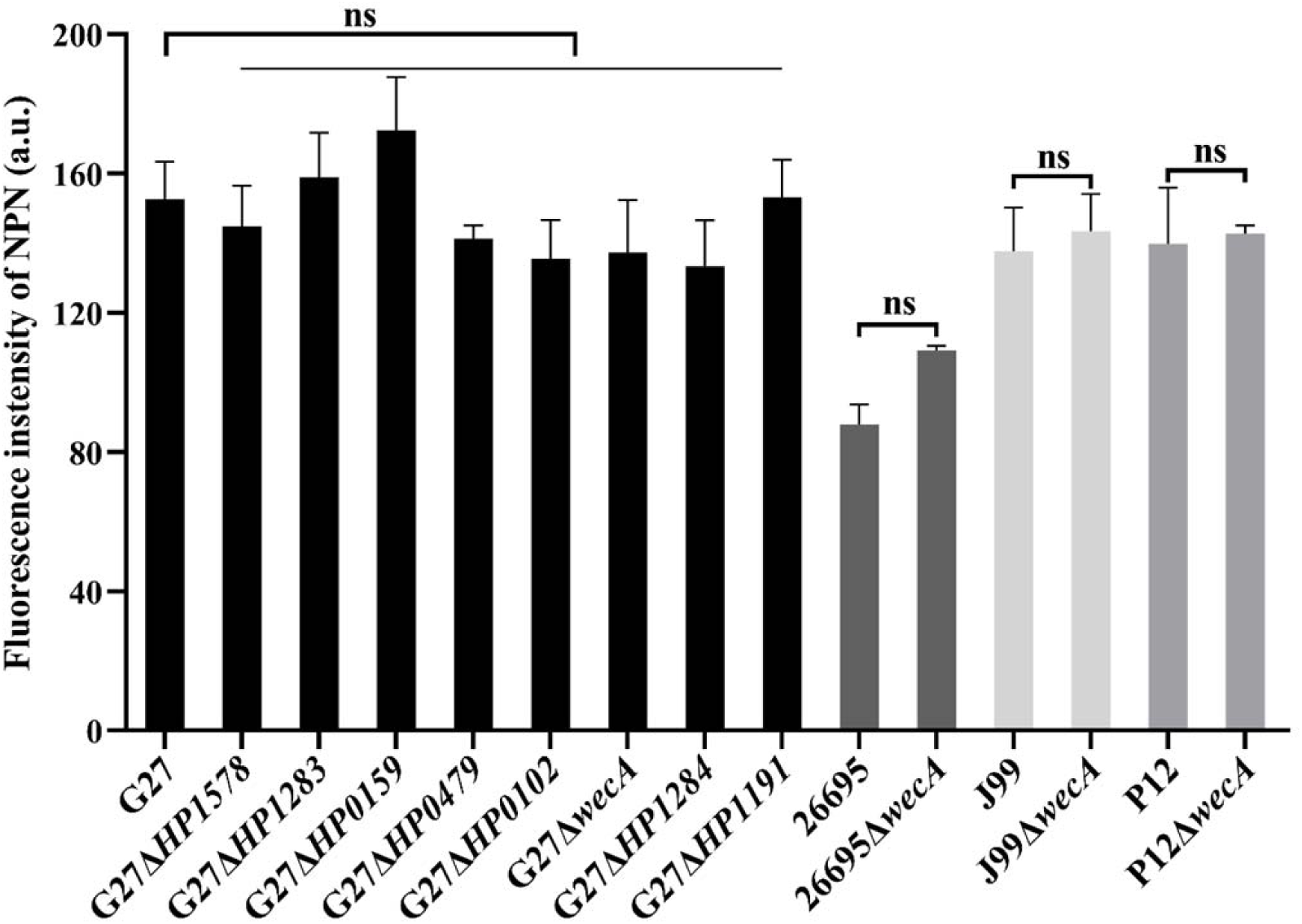
Deletion of LPS glycosyltransferase genes has no effect on the outer membrane permeability of *H. pylori*. Outer membrane permeability of *H. pylori* was measured by the influx of fluorescence probe N-Phenyl-1-naphthylamine. Fluorescent intensity is positively correlated with the permeability of outer membrane. Data from three independent experiments are presented as mean ± standard errors of the mean (SEM). ns, not significant. Student’s t-test, two-tailed.

We also investigated whether the deletion of various LPS glycosyltransferase genes had any effect on *H. pylori* susceptibility to other 6 antibiotics. In G27 strain background, the wild-type strain and all the 8 LPS mutants had similarly low MICs against clarithromycin, levofloxacin, amoxicillin, tetracycline, and metronidazole, and were all sensitive to these antibiotics. However, G27 wild-type strain was found to be resistant to rifampicin, whereas all the 8 LPS mutants were found to be sensitive to rifampicin. This was further confirmed in 26695, J99, and P12 strain background: the deletion of *wecA gene in* 26695, J99, and P12 strains did not change their susceptibility or MICs against clarithromycin, levofloxacin, amoxicillin, tetracycline, and metronidazole; however, 26695, J99, and P12 wild-type strains were all resistant to rifampicin, whereas all their corresponding Δ*wecA* mutant became rifampicin-sensitive (**Table 2**, **Fig 6**).

**Fig. 6.**
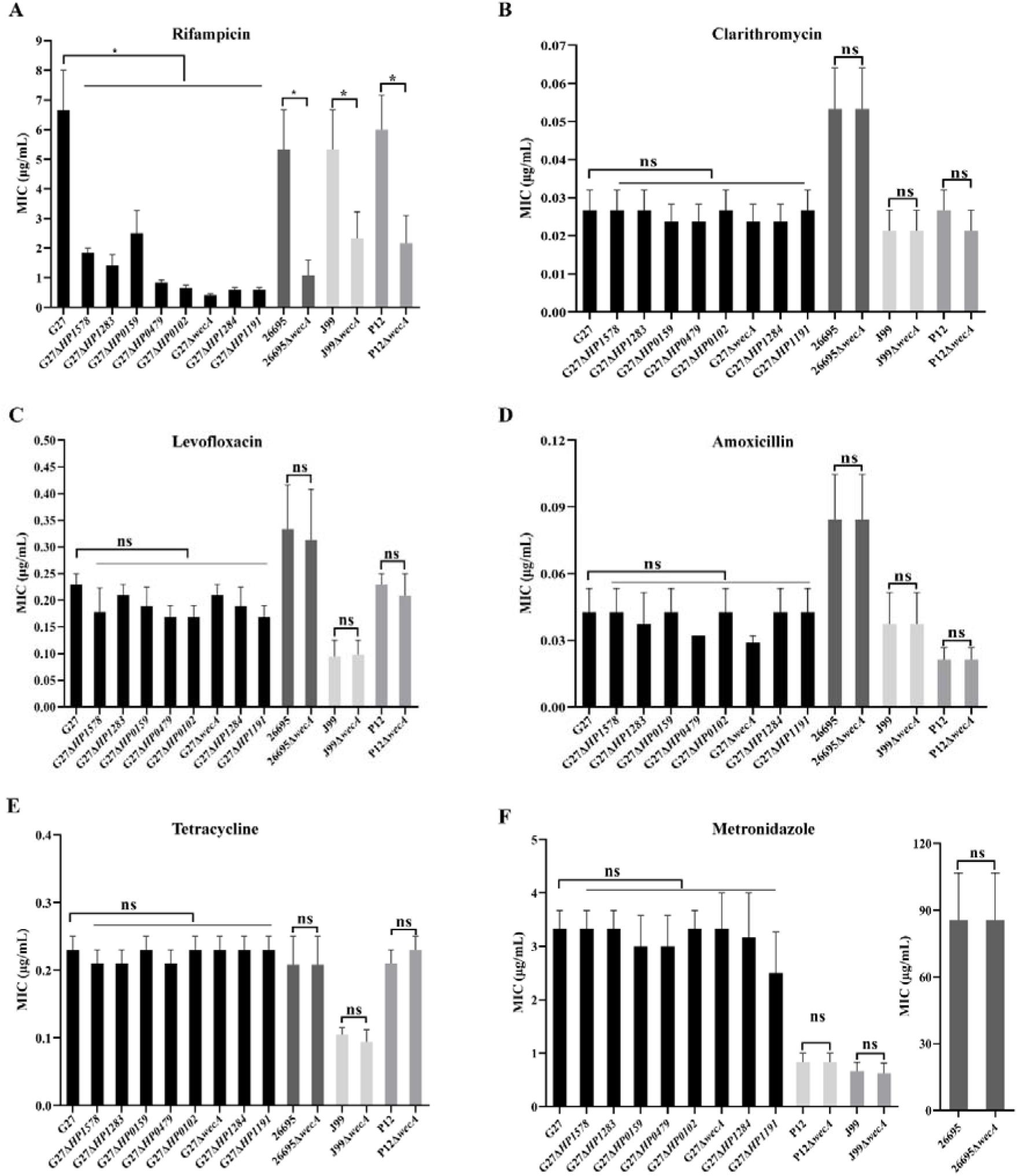
Deletion of LPS glycosyltransferase genes increases *H. pylori* susceptibility to rifampicin but not to other clinically used antibiotics. Minimum inhibitory concentrations (MICs) of wild-type *H. pylori* strains and their associated LPS mutants to rifampicin (A), clarithromycin (B), levofloxacin (C), amoxicillin (D), tetracycline (E), and metronidazole (F) tested by agar dilution assay. All values were indicated as mean with standard errors of the mean (± SEM) from three independent experiments. *, *P* < 0.05, **, *P* < 0.01, ***, *P* < 0.001; Student’s t-test, two-tailed.

## Discussion

In the present study, using a series of *H. pylori* LPS mutants, we systematically analyzed the possible roles of LPS glycosyltransferase genes in maintenance of bacterial growth, morphology, cell wall integrity, and susceptibility to antibiotics. We observed that all the LPS mutants had no defects in growth fitness. However, we observed that the deletion of LPS glycosyltransferase genes affected the spiral and rod-like morphology, especially after 48 h of culture: the cells in G27Δ*HP1578* and G27Δ*wecA* were almost completely coccoid, whereas the rod-like cell shape was still clearly observed in wild-type. Previous studies have shown that multiple proteins including cell-shape-determining proteins (Csd1, Csd2, Csd5 and Csd7) and bactofilin homolog CcmA are required to generate the helical shape of *H. pylori* cells, influencing the shape and composition of the peptidoglycan sacculus directly or indirectly.^23^ The morphology change in the LPS glycosyltransferase gene mutants could be an indirect change to the composition of the peptidoglycan sacculus. As UDP-GlcNAc is the universal substrate utilized by both the LPS pathway and the peptidoglycan pathway,^23^ it is possible that the perturbation of one pathway could influence the other. For example, as both HP1578 and WecA are the GlcNAc transferases, deletion of *HP1578* and *wecA* genes blocks the use of the intracellular GlcNAc for LPS biosynthesis, which in turn may drive synthesis of peptidoglycan, offsetting the relaxation of peptidoglycan by the multiple endopeptidases. In addition, the cell morphology change upon the deletion of LPS glycosyltransferase genes in *H. pylori* may be explained by interference of other cytoskeletal elements. For example, it has been reported that the coiled-coil-rich proteins (Ccrp), which have a similar molecular architecture as intermediate filaments are found to be associated with the helical shape of *H. pylori.*^24, 25^ In *Caulobacter crescentus*, LPS mutation was reported to interfere with the filament-like protein crescentin-mediated cell curvature, causing a disruption of normal cell morphogenesis.^26^ It should be noted that the helical cell shape of *H. pylori* plays an important role in efficient stomach colonization.^27, 28^ Previous studies reported that *HP1284*, *HP0102*, and *HP0159* mutants failed to colonize C57BL/6 mice,^12, 17, 29^ which could be partly ascribed to the morphological changes induced by the deletion of corresponding LPS glycosyltransferase genes.

The deletion of LPS glycosyltransferase genes including the G27Δ*HP1191* mutant (with LPS truncation starting from the Hep II residue) does not affect *H. pylori* outer membrane permeability. This is in sharp contrast to the markedly increased permeability phenotype observed in the deep rough LPS mutants of other Gram-negative bacteria,^30, 31^ which could be explained by the appearance of phospholipid bilayer patches in the outer membrane due to the decreased incorporation of outer membrane proteins, or the perturbation of the lateral interactions between the neighboring LPS molecules in the presence of defective LPS molecules. The crucial structure for the LPS-protein or lateral LPS-LPS interactions in these Gram-negative bacteria has been proposed to lie in the presence of the negative phosphate groups on the core Hep II residue.^32^ For example, in *E. coli* and *P. aeruginosa*, the core Hep residues are decorated with negatively charged phosphate groups, which are believed to play an important role in interacting with basic groups of the outer membrane proteins, and in enabling the neighboring LPS molecules to be cross-linked by divalent cations, and thus stabilizing the outer membrane.^33–35^ However, in *H. pylori*, the LPS are constitutively modified to have a core oligosaccharide without the presence of negatively charged phosphate groups on Hep II.^8^ Thus, the deletion of Hep II glycosyltransferase gene *HP1191* does not change the electrostatic interactions between the *H. pylori* LPS and the outer membrane proteins, or the electrostatic repulsion between the neighboring LPS molecules, which may explain the unaltered outer membrane permeability in *H. pylori* deep rough LPS mutant (G27Δ*HP1191*). Of note, Hep I glycosyltransferase gene *HP0279* deletion mutant in *H. pylori* has never been successfully constructed or identified,^17^ it may suggest that *HP0279* is essential, and that the minimal LPS structure required for viability and outer membrane permeability of *H. pylori* consists of the lipid A, KDO, and the Hep I residue.

*H. pylori* is naturally resistant to the positively charged polymyxin B, an experimental substitute for CAMPs in laboratory settings.^8^ It is well documented that the removal or “masking” of lipid A by phosphate groups is the primary mechanism involved in resistance to CAMPs.^36^ The constitutive removal of the lipid A 1’-phosphate group and 4’-phosphate group by LpxE and LpxF, respectively, reduces the negative charge of lipid A, thus rendering *H. pylori* natural resistance to polymyxin B.^10^ In the present study, we observed that the polymyxin B MICs in the G27Δ*HP1578* (lacking Lewis antigen) and G27Δ*HP1283* (lacking heptan and Lewis antigen) were comparable to that of the G27 wild-type strain. This is in line with previous reports that the variable heptan and Lewis antigen structures are not essential for establishing colonization in mouse models.^8^ Interestingly, we observed that deleting any of the conserved LPS glycosyltransferase genes including *HP0159*, *HP0479*, *HP0102*, *wecA*, *HP1284*, and *HP1191* led to a marked increase in susceptibility to polymyxin B. Of note, irrespective of parent strain background, deletion of *wecA* displayed a dramatic decrease in MIC to polymyxin B compared to wild-type. As the outer membrane permeability was unaltered among all the LPS mutants (especially the Δ*wecA* mutant), the increased susceptibility to polymyxin B might be attributed to an increased net negative charge of the mutants’ lipid A due to inefficient modification. In *H. pylori,* the lipid A-core is assembled in the cytoplasm, and translocated by MsbA to the periplasmic face, where it is constitutively modified by a five-step enzymatic pathway.^8^ It is possible that the truncated lipid A-core in the Δ*wecA* mutant may not serve as a good substrate for LpxE and LpxF, leading to an inefficient removal of the negatively charged phosphate groups, and resulting into an increased sensitivity to polymyxin B.

In the present study, we showed that none of the 8 LPS glycosyltransferase gene mutants had increased susceptibility to the commonly used 5 anti-*H. pylori* antibiotics (clarithromycin, levofloxacin, amoxicillin, tetracycline, and metronidazole) as compared to the wild-type strain. This was understandable as the outer membrane permeability was not affected among all the LPS mutants. However, we observed that deleting any of the LPS glycosyltransferase genes rendered *H. pylori* sensitive to rifampicin. Of note, rifampicin is an effective antibiotic against Gram-positive bacteria, but less effective against Gram-negative bacteria due to the impermeable LPS outer membrane.^37^ In this study, we showed that G27, 26695, J99, and P12 wild-type strains were all resistant to rifampicin, whereas deletion of *wecA* rendered them sensitive to polymyxin B. As all the LPS mutants had unaltered outer membrane permeability, and in combination with the fact that rifampicin carries one positive charge favoring interaction with LPS,^38^ the increased susceptibility to rifampicin due to the deletion of LPS glycosyltransferase genes might also be attributed to an increased net negative charge of the mutants’ outer membrane.

In conclusion, we have shown that LPS glycosyltransferase genes played an important role in the maintenance of *H. pylori*’s morphology, and the deletion of these genes resulted in significant morphological changes (coccoid, coiled “c”-shape, and irregular shapes) after 48 h growth as compared to the rod-like cell shape of the wild-type strain. Moreover, we showed that deletion of conserved LPS glycosyltransferase genes does not affect *H. pylori* outer membrane permeability, but increases bacteria sensitivity to polymyxin B and rifampicin. Of note, irrespective of parent strain background, deletion of *wecA* displayed a dramatic increase in susceptibility to polymyxin B. Interfering LPS biosynthesis through the deletion or inhibition of the key LPS glycosyltransferase genes (like the *wecA*) does not kill the bacteria but rather disable the bacteria establishing successful colonization. Thus, conserved *H. pylori* LPS glycosyltransferases, which are essential for typical helical morphology and polymyxin B resistance could be promising targets for developing novel drugs aiming at “disarming” *H. pylori*.

## Materials and methods

### Bacterial strains, plasmids, growth conditions, and oligonucleotide primers

Bacterial strains and plasmids used in this study are summarized in **Table 1**. The following *H. pylori* strains were used: G27 wild-type strains and the associated eight LPS truncated strains, 26695, J99, P12 wild-type strains and their Δ*wecA* mutant strains. *H. pylori* strains were cultured in Columbia blood agar (CBA) plates supplemented with 5% defibrinated sheep blood and 5% fetal calf serum (FCS) and incubated at 37℃ under microaerobic condition (85% N_2_: 5% H_2_: 10% CO_2_) generated by the Anoxomat Mark-II system (Mart Microbiology B.V., the Netherlands). DNA oligonucleotide primers for the identification of *wecA* deletion in 26695, J99, and P12 are named as WecA-F with sequence (5’->3’) of CACGCTATGACCGATATTAAGC and WecA-R with sequence (5’->3’) of GCTGTTCTGTTTGAGACAAG.

### Construction of *wecA* gene mutant in *H. pylori* 26695, J99, and P12

For constructing *wecA* mutant in *H. pylori* 26695 (streptomycin resistant), we used a highly efficient Xer-cise gene deletion method.^39^ Briefly, 26695 was transformed with previously constructed plasmid pWecA-AB-difH-RC and cultured in BHI blood agar plates (brain heart infusion, 5% defibrinated sheep blood and 5% FCS) supplemented with chloramphenicol. Single colonies were sub-cultured in BHI blood agar plates with streptomycin to generate clean deletion of *wecA* in 26695. Similar to the construction of *wecA* in 26695, plasmid pWecA-AB-difH-RC was used to transform strain J99 and P12 (both streptomycin sensitive) to generate replacement knockout of *wecA* with *difH-rpsl-cat-difH* cassette.

### *H. pylori* LPS microextraction for silver staining

*H. pylori* LPS was micro-extracted as previously performed. Briefly, bacterial cells with an amount of OD_600_ = 3 were harvested from CBA plates and suspended in LPS lysis buffer that composed of 2% SDS, 4% β-mercaptoethanol, 0.1% bromophenol blue, 10% glycerol, 1 M Tris-HCl (pH 6.8). Samples were heated at 100℃ for 10 minutes and then cooled. Thereafter, the samples were added with 5 μL proteinase K (20 mg/mL) and incubated in 55℃ of water overnight. The obtained LPS samples were run in 15% SDS-PAGE gels. After the gels underwent staining through three steps: oxidation, silver dyeing and color development, the LPSs were visualized thoroughly for structural analysis.

### Morphology analysis of *H. pylori* strains by gram-staining

One drop of saline was added on a clean glass slide. Bacteria colonies were transfer with a plastic loop from the surface of agar plate onto the saline drop and spread evenly. The bacterial solution was air dried and stained with Gram’s crystal violet for 1 min, and then rinsed with tap water. Subsequently, the bacteria were stained with iodine solution for 1-3 min and rinsed with tap water. The excess water was gently removed with a tissue paper. Thereafter, the bacteria were counter-stained with Carbol Fuchsin for 1 min and rinsed with tap water, and the excess water was gently removed with a tissue paper. Finally, the stained bacteria were mounted with coverslip by paraffin and visualized using Olympus Microscope (Olympus Corporation, Japan). Digital images were taken using a camera connected with the microscope via a microscope adapter.

### Growth analysis of *H. pylori* strains

The growth properties of the *H. pylori* LPS mutants was determined in brucella broth supplemented with 10% FCS (BB10). Individual 5 mL cultures of BB10 were inoculated with G27 wild-type or LPS truncated strains to give a starting concentration of bacterial solution as OD_600_ = 0.1. Cultures were incubated at 37℃ with 100 rpm shaking under microaerobic condition. The optical densities of the cultures were measured every 6 h and/or 12 h for up to 48 h.

### Outer membrane permeability assay

Fluorescent probe N-Phenyl-1-naphthylamine (NPN) (Sigma-Aldrich USA) was used to examine the outer membrane integrity of *H. pylori* wild-type and its LPS mutant strains, as described previously.^40^ Briefly, *H. pylori* strains were harvested from CBA plates in logarithmic growth stage, and then washed for three times in HEPES (pH 7.0, 5mM). The bacterial pellets were suspended with HEPES and OD_600_ value of obtained bacterial suspension was adjusted to 0.4. Afterward, the dye NPN was added to the bacterial suspension, achieving a final concentration of 10 μM. The NPN-bacteria mixture was incubated at 37℃ for 30 min, away from light. Finally, the mixture was transferred to a 96-well plate with black walls at 200 μL per well, following fluorescence measurement on a Varioskan Flash-Full Wavelength Microplate Reader (Thermo Scientific, USA) with the excitation wavelength at 350 nm and the emission wavelength of 420 nm.

### Antimicrobial susceptibility testing

The minimum inhibitory concentrations (MICs) of polymyxin B and six clinically used antibiotics including clarithromycin, metronidazole, levofloxacin, amoxicillin, tetracycline, and rifampicin to *H. pylori* wild-type and LPS mutant strains were determined by standard agar dilution method. A range of antibiotic concentrations was prepared by twofold serial dilutions of antimicrobial (ranging from 0.125 to 128 μg/mL) incorporated into BHI blood agar plates. The inoculum was delivered onto the surface of the agar plates with antibiotics by a Multipoint Inoculator (Denley, UK) to obtain approximately 1 × 10^5^ CFU per spot. After static incubation for 3-5 days at 37℃ under microaerobic condition, the MIC was defined as the lowest concentration of drug inhibiting visible *H. pylori* growth. According to the recommendation of European Committee on Antimicrobial Susceptibility Testing (EUCAST), version 10.0, 2020, resistance to clarithromycin, metronidazole, levofloxacin, amoxicillin, tetracycline, and rifampicin was defined as MIC > 0.5 mg/L, > 8 mg/L, > 1 mg/L, > 0.125 mg/L, > 1 mg/L and > 4 mg/L, respectively.^41^

### Statistical analysis

The software Graphpad Prism version 9.1.2 (SPSS Inc., Chicago, USA) was used for the statistical analyses in this study. The significance of different values from different *H. pylori* strains was determined by Student’s t-test, with a *P* value of less than 0.05.

